# Collation of indigenous and local knowledge as evidence base for herpetofauna conservation outside protected areas: Case study from an agricultural landscape in Eastern India

**DOI:** 10.1101/2021.05.26.445747

**Authors:** Deyatima Ghosh, Parthiba Basu

**Affiliations:** Department of Zoology, University of Calcutta, 35, Ballygunge Circular Road, Kolkata, 700-019

**Keywords:** Amphibian, Agricultural Intensification Gradient, Community Knowledge, Indigenous Knowledge, Eastern India, Reptile

## Abstract

Systematic appraisal of community’s knowledge as evidence for biodiversity conservation has been widely recognized. For conserving the rich biodiversity in the rural landscape outside the protected areas, it is important to document the knowledge and perception of the farming community. Although such appraisal is available for different taxa, no such systematic study is available for herpetofauna-one of the most vulnerable faunal groups. Our study attempts to document the impact of agricultural intensification on herpetofauna in an agricultural landscape through knowledge and perception appraisal of the farming community. A semi-structured questionnaire survey and validation was conducted in areas of low, medium, and high agricultural intensification. In all areas, farmers indicated an overall decrease in herpetofauna abundance. Farmers at the mid and high agricultural intensification zones reported a more significant decrease in herpetofauna sightings specifically for amphibians and snakes compared to those under low intensification regions. Farmers at low intensification area recognized significantly more herpetofauna. Farmers attributed five major threats to herpetofauna and ranked pesticide as the most significant reason, especially those in higher intensification. The majority were aware of the importance of herpetofauna as a biological pest control agent. Level of education or farming experience did not seem to have any influence on the farmers’ knowledge. Our findings integrated with other quantitative studies will facilitate future community-driven conservation in the studied agricultural landscapes.

## Introduction

Importance of systematic appraisal of the local community knowledge about the status or conservation threats to biodiversity has been widely recognized (Sutherland et al. 2004; Brook and McLachlan 2008; Chowdhury and Koike 2010; Singh et al. 2013; Braga and Schiavetti 2013; Roue et al. 2017). Use of multiple evidence that recognizes the community’s knowledge in shaping conservation programs has therefore been strongly advocated (Hunter and Brehm 2003; Sutherland 2013; Segger and Phillips 2015; Dicks et al. 2016; Smith et al. 2017). Nagoya Protocol (2014) too emphasizes the role of local and indigenous knowledge for conservation and sustainable use of biodiversity. Notwithstanding the importance of ecological studies in addressing issues concerning the status and sustainability of biodiversity, community-based information can bring a much-needed focus in improving our understanding as the strength of such knowledge lie in long exposure to series of local observations and changes (Folke et al. 2003) that conventional quantitative ecological studies often cannot capture. Such traditional ecological knowledge is the generation, accumulation, and transmission of information across generations often resulting in adaptive management of local ecological resources (Berkes Colding and Folke 2000).

Agricultural lands account for a vast expanse of the landscape beyond protected network and harbor rich biodiversity that is often overlooked. Several studies have taken this approach of appraising from the farming community about different biodiversity elements and species providing ecosystem services in agriculture (Singh and Sureja 2006; Cesard and Heri 2014; Grzywacz et al. 2014; Smith et al. 2017). However, studies with a similar approach are non-existent when it comes to farmland amphibians and reptiles or the herpetofauna.

The herpetofauna epitomizes global biodiversity meltdown (Gibbons 2000; Collins and Storfer 2003; Stuart et al. 2004; Beebee and Griffiths 2005) and is one of the most vulnerable among threatened faunal groups. Agricultural management leading to habitat loss (Fabricius et al. 2003; Scoccianti 2004; Araujo et al. 2006; Whitfield et al. 2007) and pesticide usage (Kittusamy et al. 2014; Sparling et al. 2015) is a significant driver of their decline. They are all the more vulnerable due to their cryptic habit and sensitivity to microhabitat changes (Valentine et al. 2007). Although recent studies have highlighted their ecological roles in the agricultural systems-little is known on the subject (Khatiwada et al. 2016; Teng et al. 2016).

Our study attempts to document a range of information from the farming community regarding herpetofauna-1. If the farmers notice any change in herpetofauna diversity in their immediate landscape. 2. How familiar are they with the herpetofauna at locations with different levels of agricultural intensification? 3. What threats do these local farming communities attribute to any change in herpetofauna diversity and abundance? 4. How do they value the ecological roles of the herpetofauna?

## Materials and methods

### Study site

The study was conducted in Balasore District (21° 29′ 41.9208″ N, 86° 56′ 33.5652″ E) Odisha, a state in South-Eastern India. The average altitude of the district is 19.8m. Balasore District covers an area of 3,634 km^2^. Broadly the district can be divided into three geographical regions, namely, the coastal belt, the inner alluvial plain, and the North-Western hills. It is bounded by the Midnapore district of West Bengal in its North, the Bay of Bengal in the east, Bhadrak district in the South, and Mayurbhanj and Keonjhar districts on its western side. The average temperature of the district varies from 22°C to 32°C with an annual rainfall of 1583 mm (Balasore official website).

### Agricultural Intensification Gradient

Survey was conducted from June 2015 to June 2016 in 20 villages spread across three towns: Nilgiri, Remuna, and Jaleswar (Fig 1). These three areas respectively represent areas of low, medium, and high agricultural intensification (Table 1 represents the criteria for classifying the three zones) (Chakrabarti et al. 2015). Study areas were predominantly under paddy cultivation and the survey program included only farmers.

**Figure 1.**
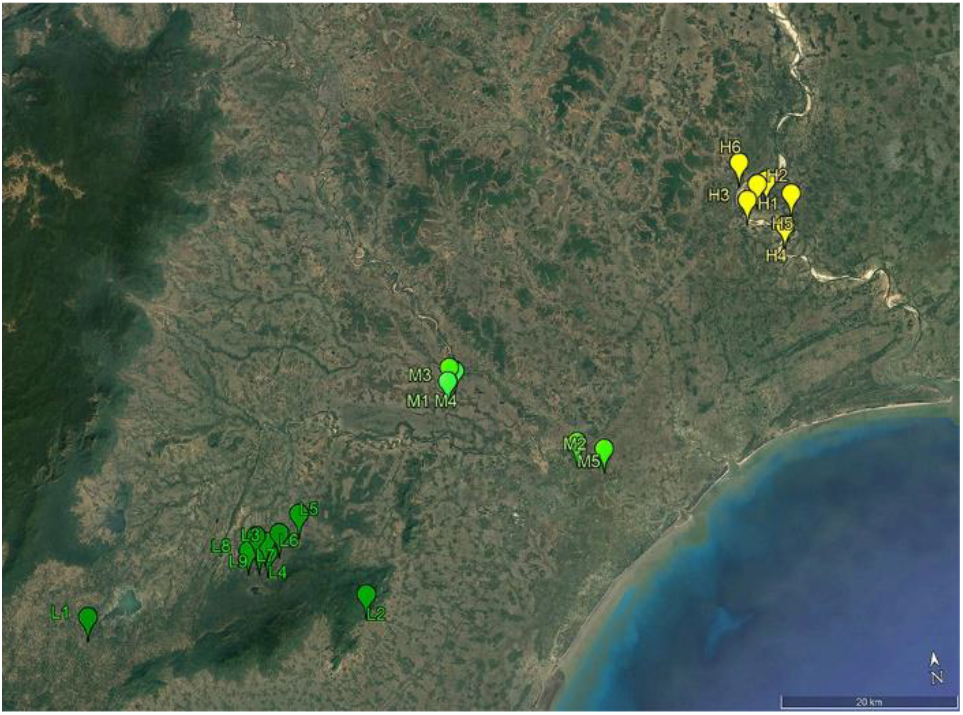
Map showing three towns along the intensification gradient where interviews were conducted

**Table 1.**
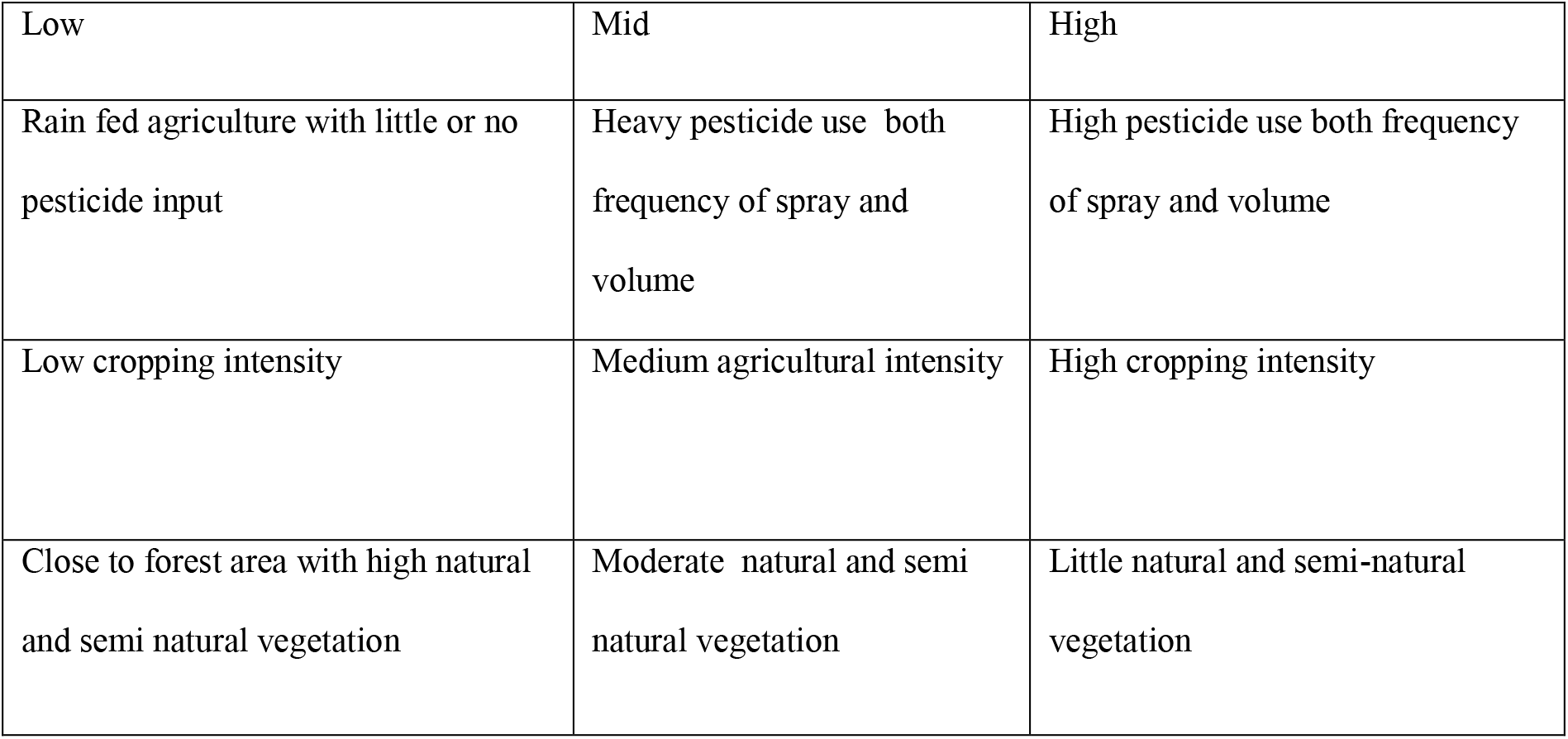
Characteristics of Low, mid and high agricultural intensification gradient.

### Survey

The overall survey followed a methodology modified after Smith et al. (2017). It was a two-step process - the first phase of individual questionnaire surveys and a second phase of validating the results obtained from the first phase.

First phase included a total of 100 individual farmers taking five farmers randomly from each of 20 villages. The survey was divided into broad themes -1. Herpetofauna abundance and sighting 2. Familiarity with locally reported herpetofauna 3. Threats for herpetofauna, 4. The reactions prevalent in the community about them and 5. The awareness within the community about their ecosystem service provisioning. Result from the interviews was grouped into specific themes and represented here as the per cent response. For recognition or familiarity test we showed a pictorial checklist of reported local herpetofauna and asked to recognize them. From this we hypothesized, if at all this recognizing ability varied across the gradient and had high reliability in the data, it could be a proxy to the difference in diversity across this gradient. Table 2 shows a list of questions used for the survey. We conducted this phase in isolation to avoid any peer influence and the entire interview was open-ended to prevent any imposing of opinions from the interviewer and allowing the participants to express their ideas and attitudes (Uyeda et al. 2014). Verbal consent was obtained from respondents before interviewing them.

**Table 2.** Questions asked to the farmers.

| Questions asked in the survey |
| --- |
| 1. How many years you are in farming? |
| 2. Do you see snakes/frogs/lizards/skinks? |
| 3. Which time of the year do you see them the most? |
| 4. Where do you see them in your field? |
| 5. Which species do you see mostly during sowing and harvesting? |
| 6. What do you do when you see them? |
| 7. Can you identify them by looking at these pictures? |
| 8. Is there any change in the number of herpetofauna over the years as observed by you |
| 9. What do you think to be the reason for their decline? |
| 10. Do you feel pesticide can be a reason for their decline |
| 11. How might pesticides affect them? |
| 12. Have you seen any evidence of pesticides affecting these animals? |
| 13. Do you think they are useful in agricultural lands? |
| 14. How do you think they might be helping in agriculture |
| 15. How do you think they help in pest regulation? |
| 16. What do you feel about these animals when you see them? |

Results from the first phase were further validated by preparing statements. Table 3 provides a sample of the statements we prepared. The validation was done in two separate groups in each village. Each validating group comprised of five new farmers who did not participate in the first questionnaire survey and had no prior knowledge about the questionnaire. Thus, a total of 40 groups (two groups in each village) making a total of 200 individuals participated in this second phase. Interviews were conducted by two people (DG and a member of the community) in local Odia language. A total of 300 farmers participated in this study. During this validation, farmers discussed each statement that was readout. Each such statement was either accepted, rejected, or modified by the participating group. This led to a set of finally agreed upon consensus statement.

**Table 3.**
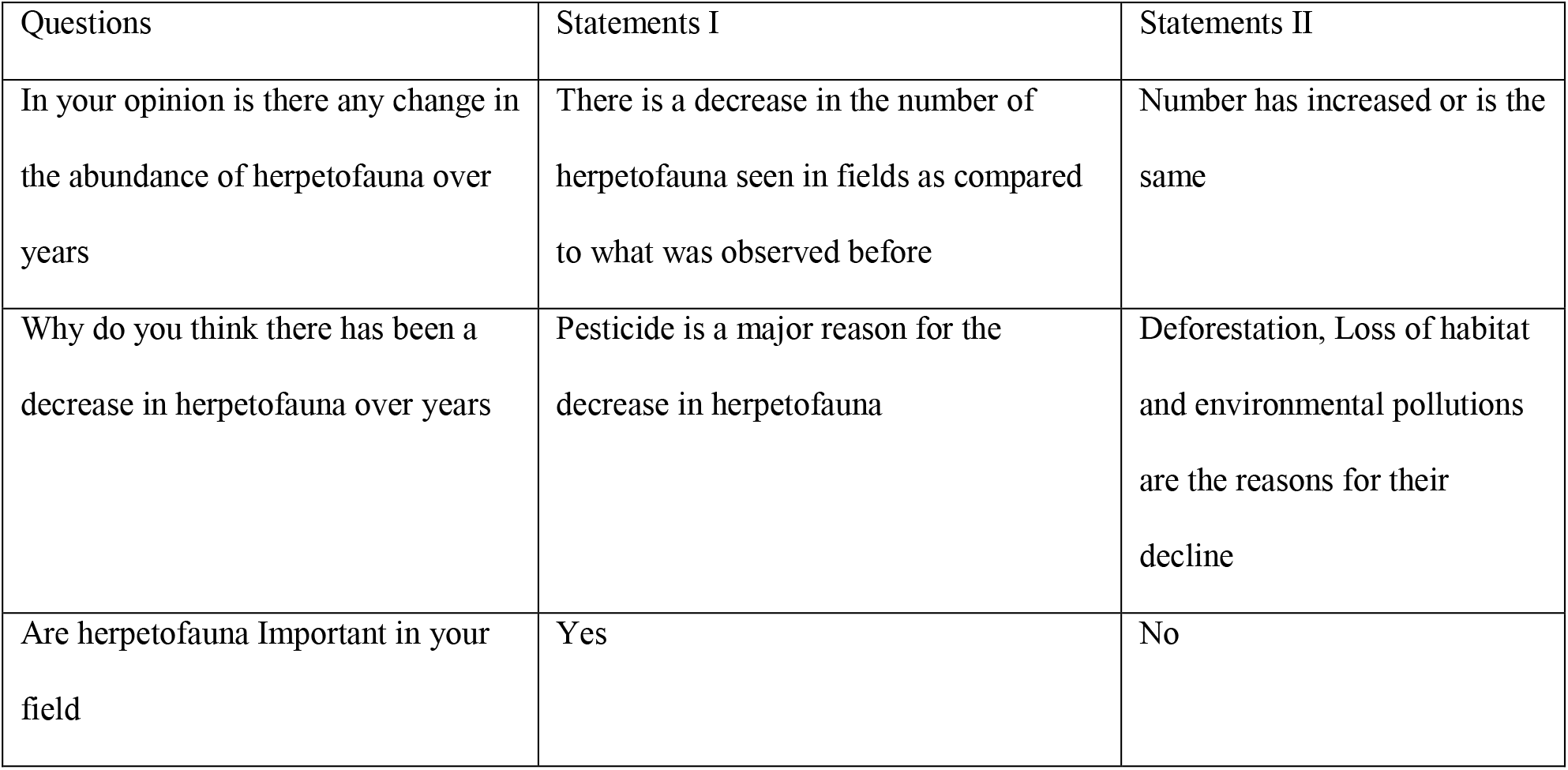
Example of few statements made from the individual questionnaire survey for the validation process.

Results from the validation phase were used to testify the findings from the phase I to enhance the quality of data and make them more conclusive. Gathering community data often suffers from lack of reliability, hence we assessed the inter-reliability (McHugh 2012) of the data obtained from phase 1 within each village (Fig. 2) and between the three groups (final checklist from phase 1 based on degree of agreement and from two validation groups; See Supplementary Information, SI1) where ever appropriate.

**Figure 2.**
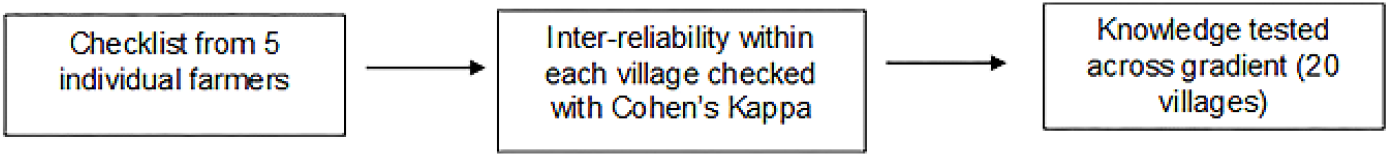
Schematic representation of performing inter-reliability with data on recognized herpetofauna in each village

## Statistical Analysis

### Effect of agricultural intensification on herpetofauna

Farmers in the individual survey were asked if they witnessed any change in herpetofauna sightings in their fields over the past few years. Chi-square test was performed to check if there existed any difference in the number of farmers across the gradient who voted for a decrease and whether it varied for any specific herpetofauna groups i.e. for snakes, lizards, amphibians, or skinks. Marascuilo post hoc test was applied to the significant chi-square results to know exactly which node of intensification made a significant impact in voting and for which specific groups.

### Community Concern for herpetofauna decline

Farmers agreeing to a decrease in herpetofauna were further questioned about the reasons they believed were responsible for their decline. These questions were open-ended and were non-suggestive. For the drivers voted by a maximum number of respondents, we checked if there was a difference in the number for that opinion across the gradient and also used results from the validation groups to get more accurate information.

### Inter reliability in data for testing their knowledge

We prepared a colored pictorial checklist of 46 herpetofauna (Table 4) that are reported from Odisha and are also available in the agricultural landscape consulting available reports (Sanyal 1993; Sarkar 1993; Daniel 2002; Smith 2003; Daniels 2005; Whitaker and Captain 2016). The colored pictorial checklist was shown to the farmers and was asked to recognize those they have seen in their agricultural lands. Farmers who were in agreement of seeing particular herpetofauna in their lands were further questioned for their local names.

**Table 4.** List of herpetofauna used for the survey program.

|  | Common Names | Scientific Name | Family |
| --- | --- | --- | --- |
| 1 | Worm Snake | *Indotyphlops braminus* (Daudin, 1803) | Typhlopidae |
| 2 | Common Sand Boa | *Gongylophis conicus* (Schneider, 1801) | Boidae |
| 3 | Red Sand Boa | *Eryx johnii* (Russell, 1801) |  |
| 4 | Indian Python | *Python molurus* (Linnaeus, 1758) |  |
| 5 | Copper-Headed Trinket Snake | *Coelognathus radiatus* (Boie, 1827) | Colubridae |
| 6 | Common Trinket Snake | *Coelognathus helenus* (Daudin, 1803) |  |
| 7 | Oriental Rat Snake | *Ptyas mucosa* (Linnaeus, 1758) |  |
| 8 | Banded Racer | *Argyrogena fasciolata* (Shaw, 1802) |  |
| 9 | Russell's Kukri Snake | *Oligodon taeniolatus* (Jerdon, 1853) |  |
| 10 | Common Kukri Snake | *Oligodon arnensis* (Shaw, 1802) |  |
| 11 | Chequered Water Snake | *Fowlea piscator* (Schneider, 1799) |  |
| 12 | Striped Keel Back | *Amphiesma stolatum* (Linnaeus, 1758) |  |
| 13 | Vine Snake | *Ahaetulla nasuta* (Lacépède, 1789) |  |
| 14 | Brown vine snake | *Ahaetulla pulverulenta* (Duméril, Bibron & Duméril, 1854) |  |
| 15 | Common Wolf snake | *Lycodon aulicus* (Linnaeus, 1758) |  |
| 16 | Common Bronze Back Tree Snake | *Dendrelaphis tristis* (Daudin, 1803) |  |
| 17 | Ornate Flying Snake | *Chrysopelea ornata* (Shaw, 1802) |  |
| 18 | Common Cat Snake | *Boiga trigonata* (Schneider, 1802) |  |
| 19 | Forsten's Cat Snake | *Boiga forsteni* (Duméril, Bibron & Duméril, 1854) |  |
| 20 | Green Keel Back | *Rhabdophis plumbicolor* (Cantor, 1839) |  |
| 21 | Monocled Cobra | *Naja kaouthia* (Lesson, 1831) | Elapidae |
| 22 | Spectacled Cobra | *Naja naja* (Linnaeus, 1758) |  |
| 23 | King Cobra | *Ophiophagus hannah* (Cantor, 1836) |  |
| 24 | Common Krait | *Bungarus caeruleus* (Schneider, 1801) |  |
| 25 | Banded Krait | *Bungarus fasciatus* (Schneider, 1801) |  |
| 26 | Bamboo Pit Viper | *Trimeresurus gramineus* (Shaw, 1802) | Viperidae |
| 27 | Russell's Viper | *Daboia russelii* (Shaw & Nodder, 1797) |  |
| 28 | Saw Scaled Viper | *Echis carinatus* (Schneider, 1801) |  |
| 29 | Bronze Grass Skink | *Eutropis macularia* (Blyth, 1853) | Skincidae |
| 30 | White-Spotted Supple Skink | *Lygosoma albopunctatum* (Gray, 1846) |  |
| 31 | Common snake skink | *Lygosoma punctate* (Gmelin, 1799) |  |
| 32 | Oriental Garden Lizard | *Calotes versicolor* (Daudin, 1802) | Agamidae |
| 33 | South Indian Rock Agama | *Psammophilus dorsalis* (Gray, 1831) |  |
| 34 | Chameleon | *Chamaeleo zeylanicus* (Laurenti, 1768) | Chamaeleonidae |
| 35 | Water Monitor Lizard | *Varanus salvator* (Laurenti, 1768) | Varanidae |
| 36 | Common Indian Monitor | *Varanus bengalensis* (Daudin, 1802) |  |
| 37 | Turtle | *Melanochelys trijuga* (Schweigger, 1812) | Testudines |
| 38 | Common Indian Toad | *Duttaphrynus melanostictus* (Schneider, 1799) | Bufonidae |
| 39 | Common Tree Frog | *Polypedates leucomystax* (Gravenhorst, 1829) | Rhacophoridae |
| 40 | Indian Bull Frog | *Hoplobatrachus tigerinus* (Daudin, 1803) | Dicroglossidae |
| 41 | Indian Bull Frog (Morph) | *Hoplobatrachus tigerinus* (Daudin, 1803) |  |
| 42 | Indian Skipping Frog | *Euphlyctis cyanophlyctis* (Schneider, 1799) |  |
| 43 | Fejervarya sp. 1 | *Fejervarya* sp. |  |
| 44 | Fejervarya sp. 2 | *Fejervarya* sp. |  |
| 45 | Theobald's Reed Frog | *Hylarana tytleri* (Theobald, 1868) | Ranidae |
| 46 | Balloon Frog | *Uperodon globulosus* (Gunther, 1864) | Microhylidae |

We checked for the inter reliability in data obtained from the five farmers for each village using Cohen’s kappa (Fig. 1) and performed a Kruskal Wallis test followed by Dunn’s test to check whether any difference existed in the herpetofauna recognized across the gradient.

### Level of formal education and farming experience affecting their knowledge

We speculated farmers’ recognizing ability could be dependent upon either formal education or on their farming experience. Hence, we performed two separate Kruskal-Wallis tests to check if farming experience and formal education varied across the gradient. Subsequently, we applied two Generalized Linear Models to check if farming experience or qualification had any effect on recognizing ability of the farmers across the gradient.

### Awareness about herpetofaunal role within the community

To assess the state of awareness in the community, farmers were asked whether they believed herpetofauna was useful and to those who approved of their role in agricultural fields we asked them in what ways did they contribute to agriculture. Results were pooled together from the farmers across all the three gradients and are represented graphically in the result section.

All statistical analyses were performed using R (version 3.2.3.) packages “Vegan”, “dplyr” maintaining the assumptions that our data required.

## Result

### Status of herpetofauna in agricultural lands

The number of opinions for a decrease in herpetofauna sighting (Fig. 3) was significantly different between the three zones (Chi-square: χ^2^= 33.344, df = 2, p =.000). Marascuilo test showed (Table 5) there was a significant difference in the decrease in herpetofauna sightings between low and mid and between low and high intensification zones but not between mid and high zones.

**Figure 3.**
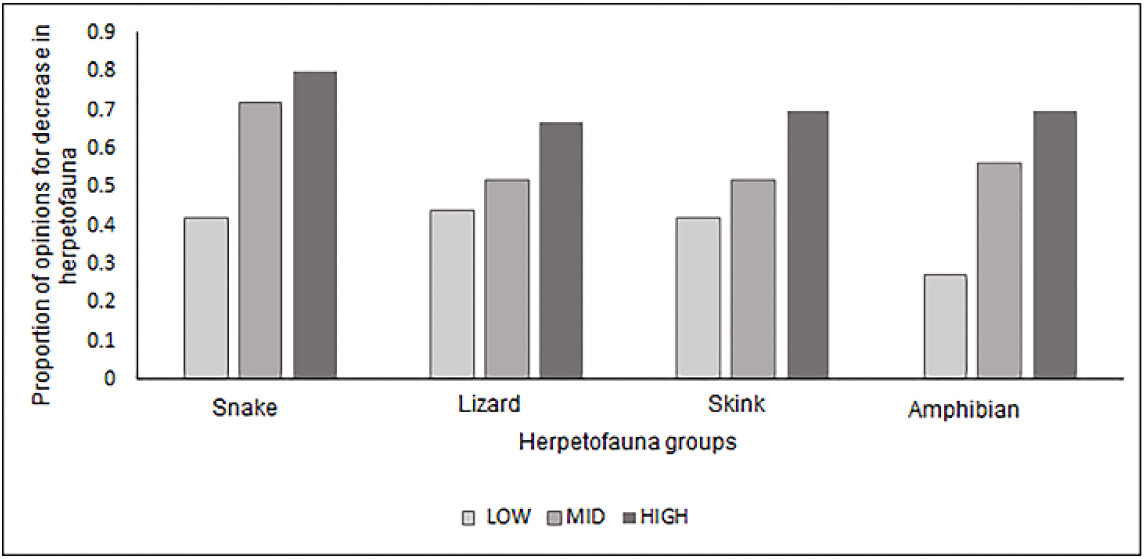
Proportion of farmers voting for a decrease of herpetofauna across gradient

**Table 5.** Marascuilo test statistics for all herpetofaunal groups.

| Difference | Proportion Difference | Critical values | Significance |
| --- | --- | --- | --- |
| Low –Mid | 0.2 | 0.1 | Yes |
| Mid- High | 0.1 | 0.2 | No |
| Low-High | 0.3 | 0.1 | Yes |

Validation results showed the same trend as well, in low intensification zones, only 6 groups out of 18 (33%), in mid intensification zones 7 out of 10 groups (70%) and 8 out of 12 (66%) groups in high intensification zones agreed upon a decrease in herpetofauna abundance.

### Which group is more vulnerable?

For individual groups of herpetofauna, significant number of farmers reported a decrease in amphibians (χ^2^= 33.344, df = 2, p = 0.001) and snakes (χ^2^= 12.494, df = 2, p = 0.001) but not for lizards (χ^2^= 4.3091, df = 2, p = 0.11) and skinks (χ^2^= 5.589, df = 2, p = 0.06). Support for this view again significantly differed between low and mid and between low and high intensification zones as observed from Marascuilo test (Table 6 & 7).

**Table 6.**
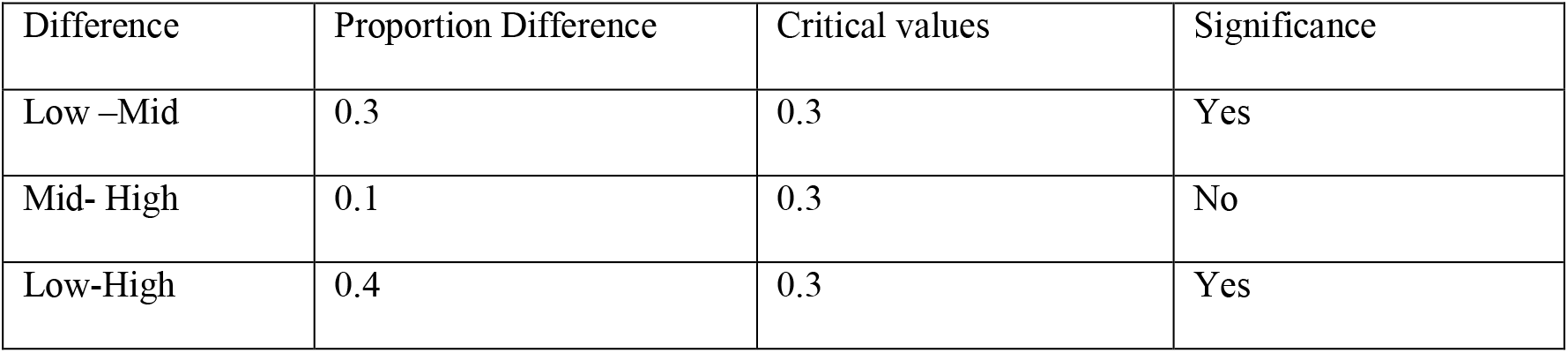
Marascuilo Test Statistics for Amphibians.

**Table 7.**
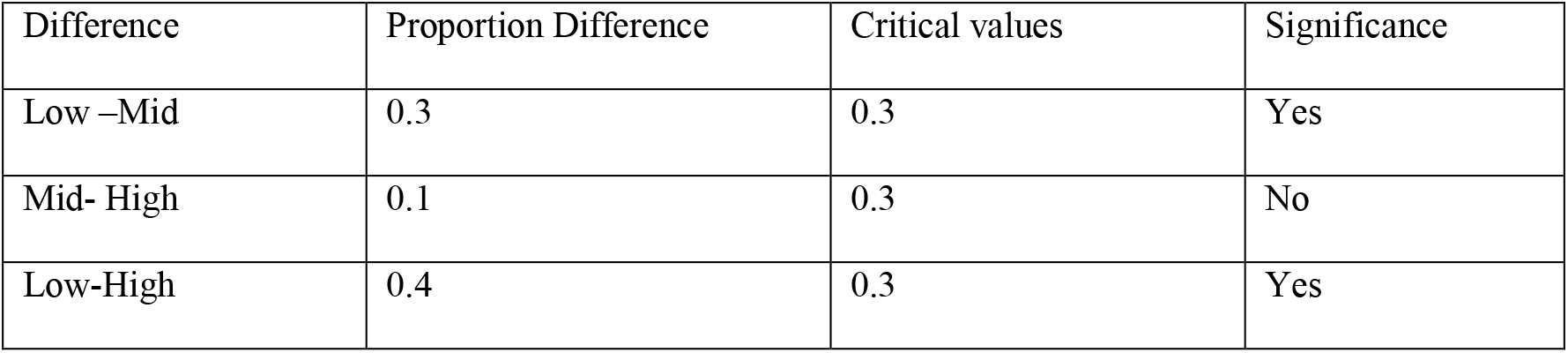
Marascuilo Test Statistics for Snakes.

Thus communities under mid and high agricultural intensification reported a more significant decrease in herpetofauna sightings specifically for amphibians and snakes compared to those under low intensification regions.

### Threats to herpetofauna: Concern within the farming community

Analyzing the results from the interviews we grouped the threats attributed by the framers in to 4 major causes e.g., deforestation, environmental pollution, deliberate killing (conflict), and pesticides application. Results are represented in figure 4. Of these, pesticide was voted to be most threatening. A similar opinion was expressed by the validating groups as well, where 36 out of the 40 surveyed groups (90%) stated pesticides to be a major threat. The number of respondents who addressed pesticide to be a threat was further compared across the gradient, and maximum farmers (73.33%) in the high intensity and mid-intensity (80%) agricultural zones viewed pesticide as one of the most crucial cause for the decrease in herpetofauna over years as compared to only 20% votes from low intensification zones (Fig. 5).

**Figure 4.**
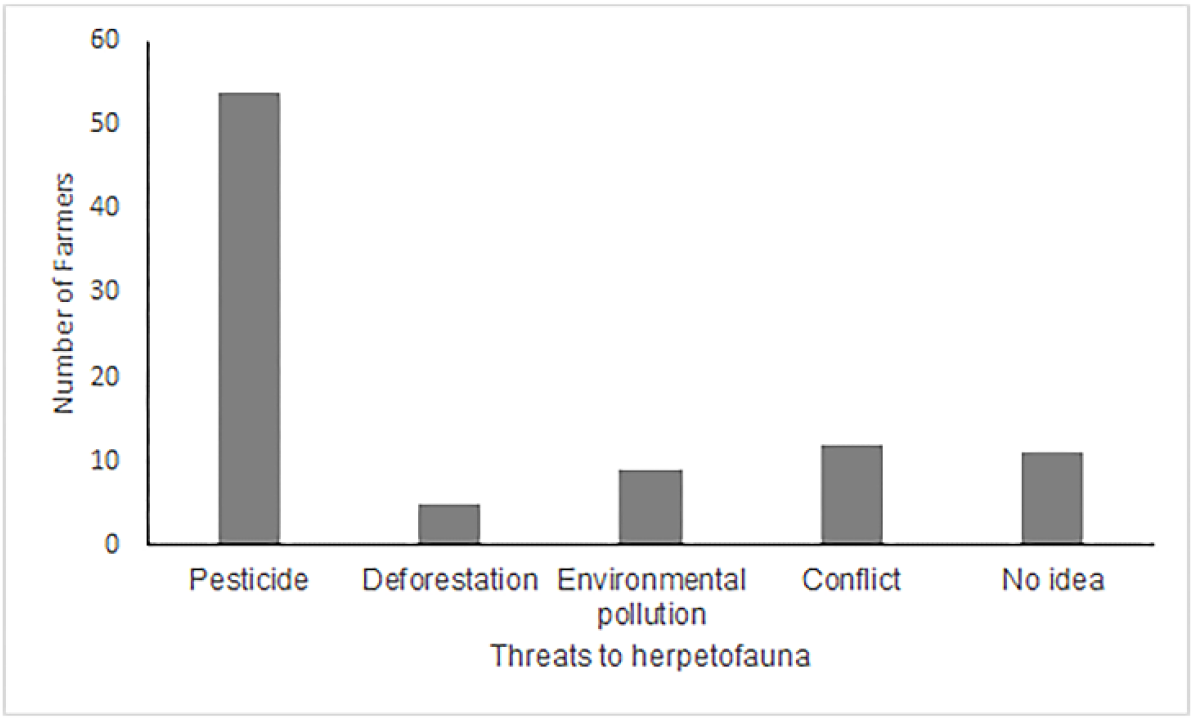
Percentage of farmers voting for deforestation, pesticide, climate/environmental pollution, conflict and pesticide as reasons for herpetofaunal decrease

**Figure 5.**
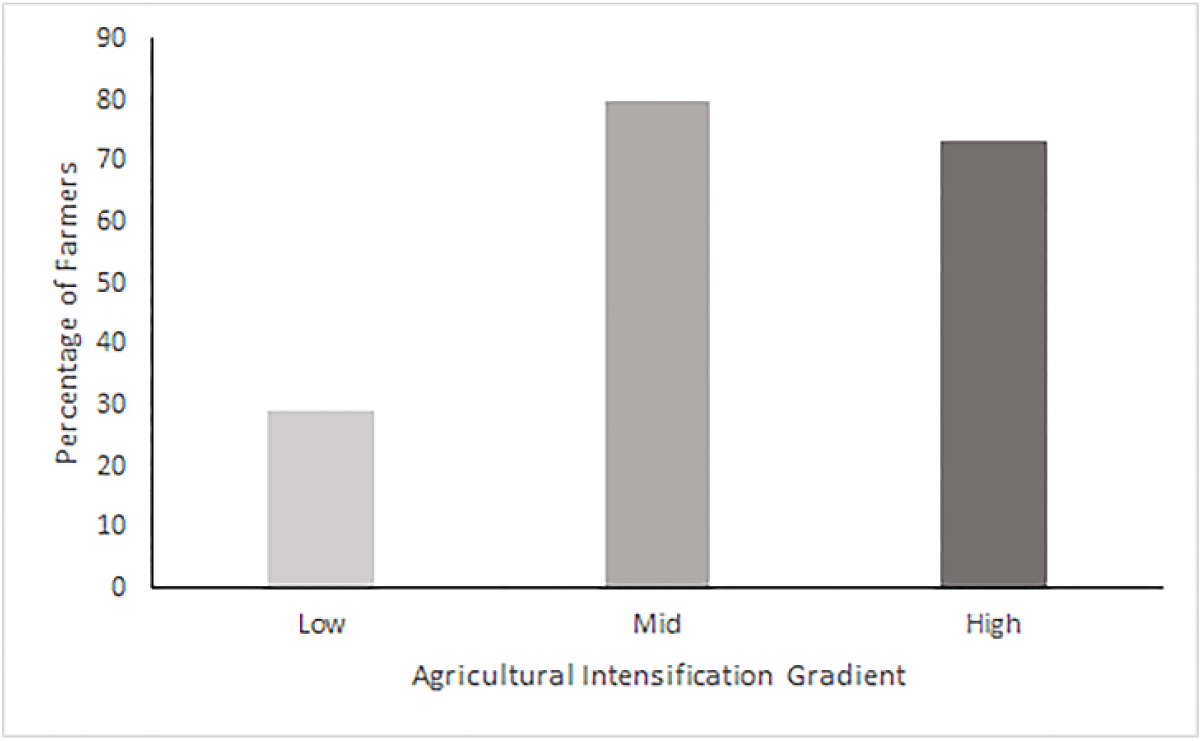
Percentage of farmers accepting pesticide as a cause for herpetofauna decrease

82% of all farmers could explain why pesticide is a direct threat for herpetofauna, especially the frogs. They also mentioned animals dying because of pesticide contamination of the water bodies in agricultural fields. Farmers were aware of pesticide contamination through the food chain. Quote from a farmer’s statement – ‘pesticide would kill the pests, frogs and lizards would feed on them which were in turn fed upon by snakes ‘. From validation data, 28 groups out of a total of 40 were in agreement with the above statements and 27 groups claimed to know this from direct field observation. From both the surveys pesticides such as Thimet, Dimecrone, Ustad (Cypermethrine) were designated as most lethal.

### Inter-reliability in data for recognizing herpetofauna

We found a high degree of agreement within the data for each village that ranged from 82% to 100%. Table 8 shows the list of Cohen’s kappa values for 20 villages. As this analysis shows, for each village, there was high inter-reliability in the data (above 64% to 100%). The two validation groups for each village (S1 Table provides the list of Cohen’s kappa for all villages) also showed high inter-reliability with the checklist prepared for each village from phase 1 hence allows for inferential statistical analysis.

**Table 8.**
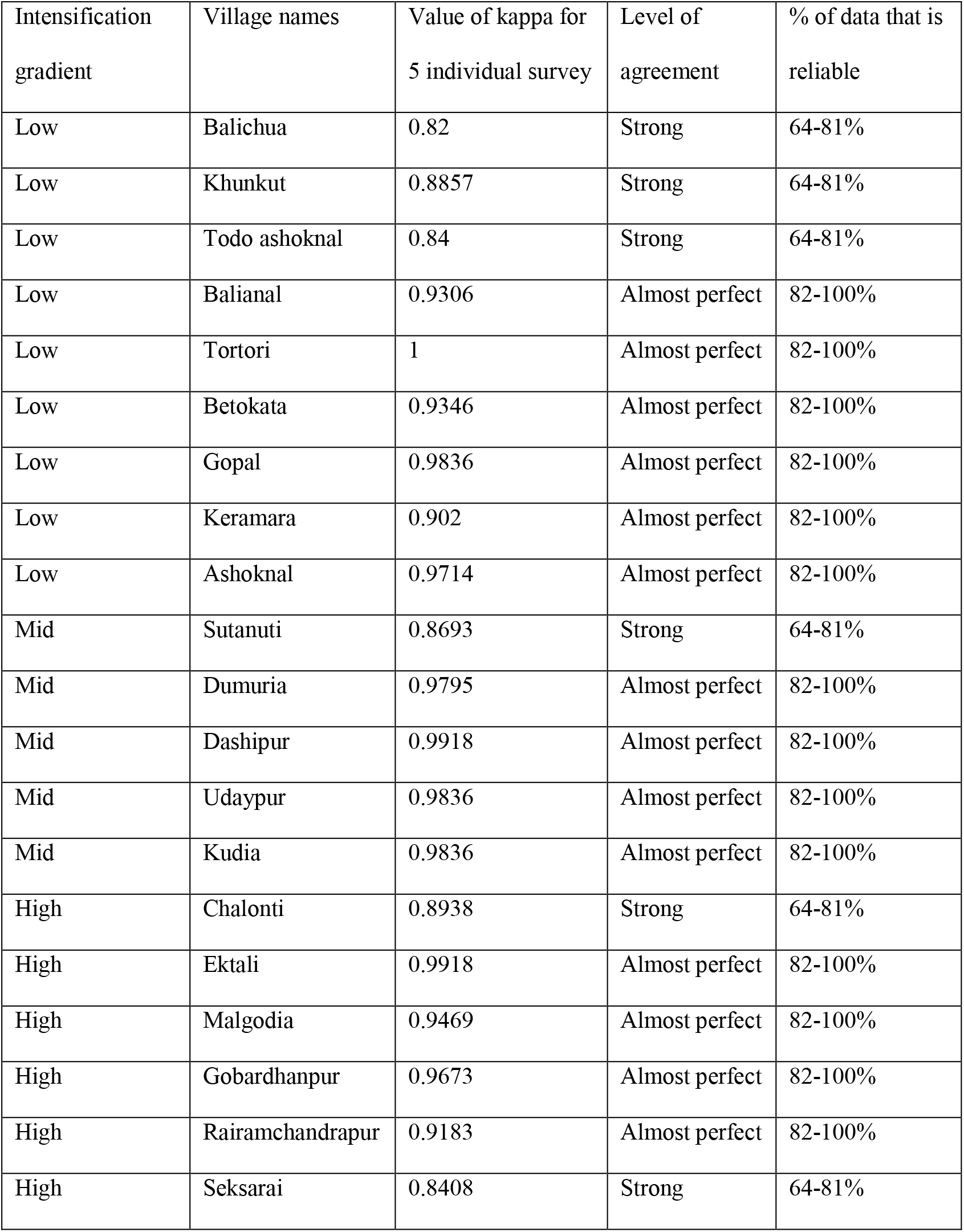
List of values for inter-reliability test for recognizing herpetofauna for each village.

### Knowledge difference across the agricultural intensification gradient

There was a significant knowledge gap among farmers, as the number of recognized herpetofauna varied significantly between low, mid, and high intensification zones (Kruskal-Wallis chi-squared = 30.129, df = 2, p=0.000). Dunn’s test showed this difference was significant between low and high (p=0.000) and low and mid (p =0.000). However, there was no significant difference between mid and high intensification zones (p = 0.6).

### Role of formal education and farming experience for recognizing herpetofauna

Formal education (Kruskal-Wallis chi-squared = 5.1108, df = 2, p-value = 0.07766) and farming experience (Kruskal-Wallis chi-squared = 0.46228, df = 2, p-value = 0.7936) did not vary across the gradient. Number of herpetofauna recognized was independent of farming experience (p = 0.238). Interestingly, formal education had significant negative effect on herpetofauna recognised (p = 0.011).

### Awareness within the farming communities

A majority of farmers were aware of the herpetofauna service provisioning in the agricultural system as 78 % of respondents across all levels of agricultural intensification agreed that herpetofauna is useful to their agriculture (Fig. 6). Further, in support of their opinions, they reported lizards, frogs, and skinks feeding on the pests and thus helping them to reduce the amount of pesticide. Small snakes, especially water snake (Checkered keelback) was mentioned to feed on the insect pests. Snakes were also reported to regulate rats, a major pest of paddy. 19 validation group supported the statement that herpetofauna feed on insect pests and nine groups were in favor of the role of snakes in controlling rats. Contradictory to this, two farmers also mentioned the negative effects of snakes in agricultural fields for digging burrows in the levees causing water drainage from agricultural lands and impairing cultivation.

**Figure 6.**
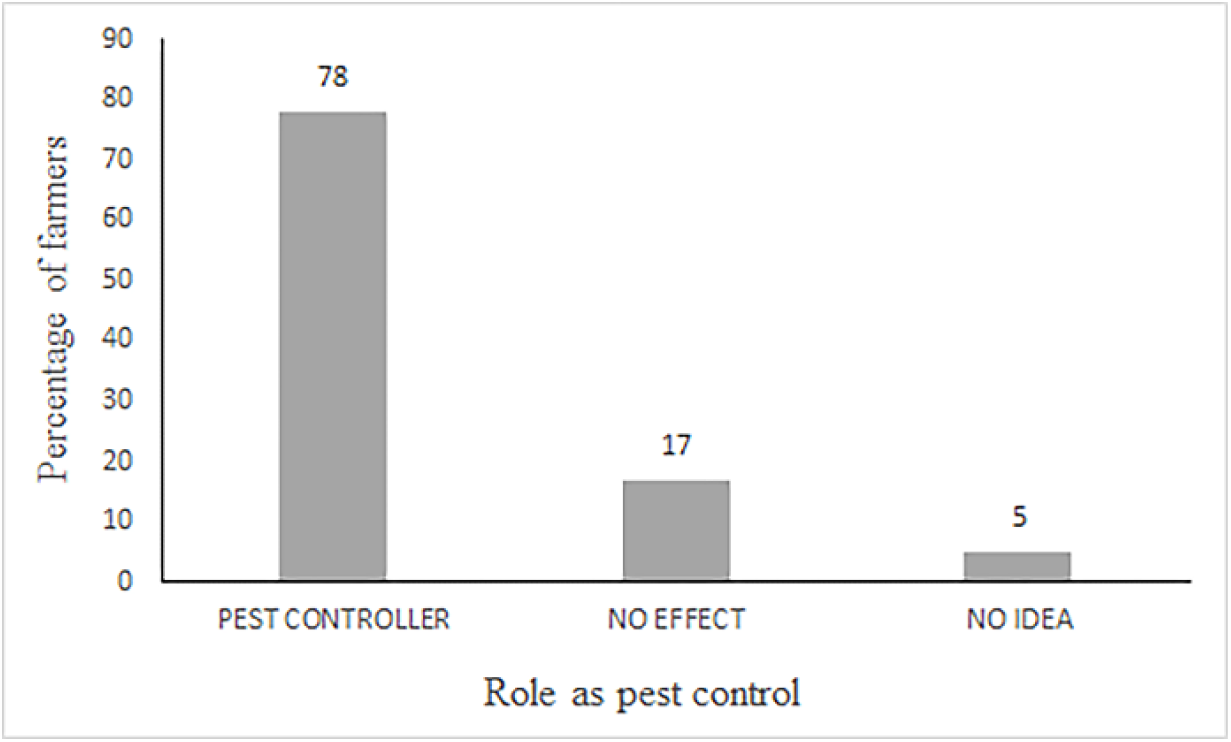
Farmers’ awareness about herpetofauna as a pest regulator

### Community reaction towards herpetofauna

As we reviewed the kinds of reactions prevalent within the farming community about herpetofauna, the most dominant attitude was that of fear from snakes followed by religious faith. Killing is also prevalent although most agreed to kill snakes that are harmful to them. However, a notable percentage of farmers seemed to be involved in indiscriminate killing as well (Fig. 7).

**Figure 7.**
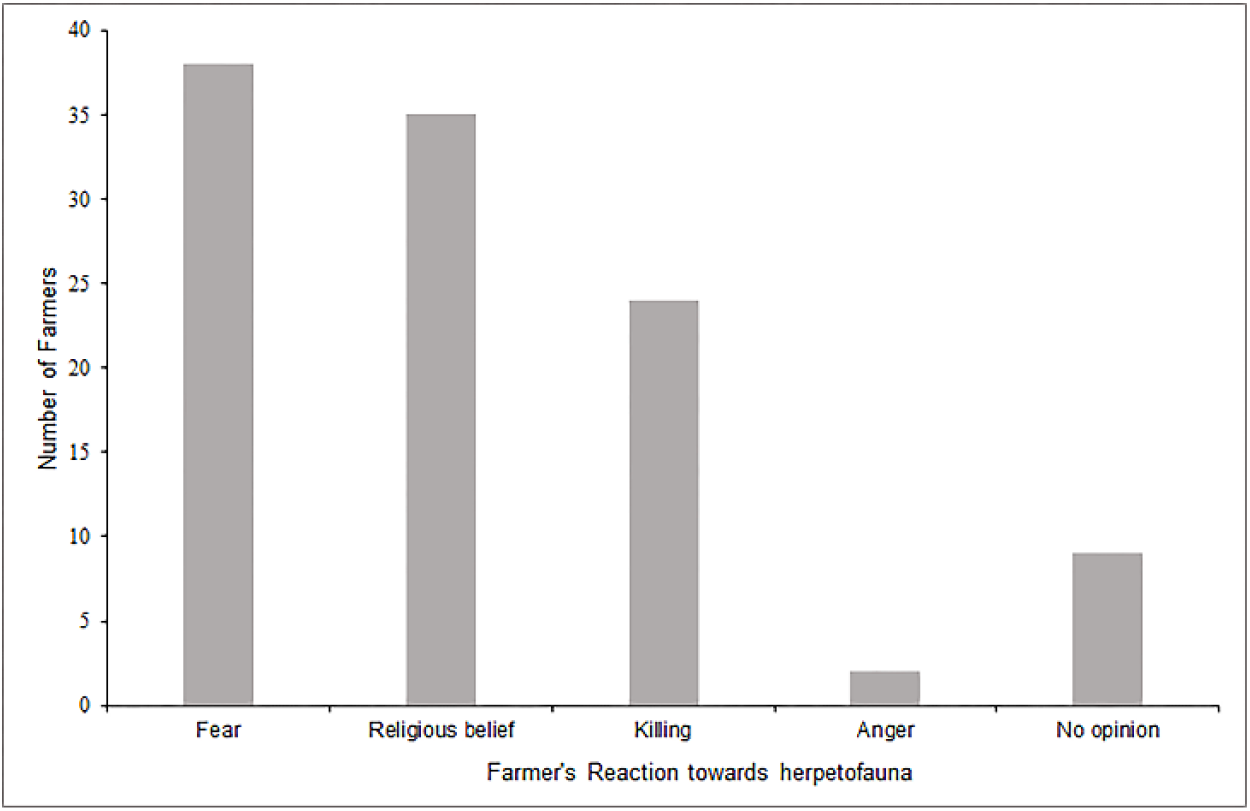
Reaction of farmers towards herpetofauna

## Discussion

Local indigenous knowledge has been identified to be a valuable source of information. Accuracy of local species knowledge is reported to be more by farmers than academic zoologists (Ulicsni et al. 2018) which occur through years of observing the environment. According to our knowledge this is the first report on assessing herpetofaunal status in an agricultural landscape from eastern India based on local ecological community knowledge. Farmers showed a distinct difference in knowledge, opinions, and perceptions of herpetofauna between the low and the higher agricultural intensification zones. Respondents in high and medium cropping intensity areas recognizing the lesser number of herpetofauna compared to the respondents in low cropping intensity agricultural areas could be either due to actual loss of diversity in the higher intensification zones or could be for some bias associated with farming exposure or due to differences in their level of education. However, no dependency on farming exposure prompt towards the diversity reported being indeed low in high cropping intensity areas compared to the areas at lower levels of intensification. Negative effect of formal education on recognizing a herpetofauna further shows the familiarity with local herpetofauna was based on practical knowledge and encounters that they have gained over years of coexistence with the taxa. Local knowledge suffers from lack of validation (Nadasdy 2005) and hence not accepted by academic scientists who are not familiar with the traditional knowledge (Gilchrist and Mallory 2007). High inter reliability in our data therefore makes the results more reliable and acceptable scientifically. Hence the difference in knowledge reported in our study is actually a true representation of the diversity status rather than occurring by chance. Familiarity with local species reflects the awareness and involvement with their immediate environment and are of high value as these communities are the custodians and play significant role in conserving and restoring the local biodiversity (Ribot 2004). Ethnozoology has wider application in science from monitoring to population biology, conservation biology and biodiversity assessment (Diamond and Bishop 1999; Colding and Folke 2001). Such information could therefore serve as baseline for conserving herpetofauna.

A few ecological studies have highlighted the effect of agricultural intensification on the diversity of herpetofauna especially amphibians (Joly et al. 2001; Beja and Alcazar 2003). Similarly, Bohm et al. (2013) has also identified agriculture as a threat to 74% of the reptilian species. These are supportive of our findings where more farmers from higher intensification areas reported a significant decrease in amphibians and reptiles in their farmlands compared to farmers in low intensification areas.

Pesticide has long been implicated as the most serious threat for the herpetofauna (Campbell and Campbell 2002; Hamer et al. 2004; Mann et al. 2009). Farmers seem to have the same opinion. This observation was affirmed more by farmers at high and mid intensification zones compared to the low zone is understandable. The intensive agriculture is overwhelmingly pesticide dependent and its impact on herpetofauna is definitely stronger in the high and the mid-level of intensification. Farmers also reported dead snakes and frogs near the agricultural field following pesticide application. Respondents felt that pesticide was not only a concern for herpetofauna but also fishes, crabs, earthworms, or any other useful fauna in agricultural landscapes.

One of the major challenges facing the conservation of herpetofauna in the agricultural landscape that our interaction with the farmers brought up was fear and antipathy. Some farmers expressed strong opposition to any proposal for saving snakes. Such fears are often (Ceriaco 2012) and could be due to aversion (Bjerke, Kaltenborn and Thrane 2001), sometimes stemmed from cultural issues or even emotional reactions (Knight 2008). This is very similar to the perceptions that Whittekar and Shine (2000) described in their questionnaire survey based on venomous snakes and human conflict in an agroecosystem. They accounted one-third of half of the times a snake when approached was killed due to their defensive strategies being misinterpreted for attacking behavior. Two contrasting reactions were documented within the farming communities – one was to spare venomous snakes revering to some religious belief while the other was to kill them due to threat to farmers working in fields. The former is a result of fear and myths and has positive implications in the conservation of particular species (Khan Menon and Bawa 1997; Berkes 1999; Devereux 2000; Swamy Kumar and Sundarapandian 2003). A noticeable percentage of farmers also concluded of killing snakes irrespective of whether they are harmless or venomous, and this was out of sheer fear (Somaweera et al. 2010; Ceriaco 2012) and often lack of knowledge (Ceriaco 2012). Nevertheless, despite overall antipathy about these animals, farmers in our study regions were aware of the benefits of herpetofauna in an agricultural system. In our study some farmers even put forth the idea of maintaining herpetofauna in agricultural land for efficient biological pest control. This has been shown in ecological studies by Teng et al. (2016), Fang et al. (2019) and Khatiwada et al. (2016). Surprisingly farmers mentioning Checkered keelback capable of pest regulation also has been proved in a study by Hossain (2016) where the study showed their diet consisting majorly of arthropods. Though there were mixed opinions and reactions, yet the fact that the majority of farmers were aware of the ecosystem service rendered by herpetofauna is a good starting point to raise a concern about these taxa.

This study evidently shows the importance of local knowledge in interpreting the deteriorating effect of agricultural intensification on farmland herpetofauna. Such local knowledge is a perfect mixture of scientific and practical evidence (Olsson and Folke 2001) and can provide baseline information and bridge the gap between conservation science and local knowledge and improve the efficiency of scientific conservation designs for concerned taxa. This study is believed to pave the path for better cooperation between academic science, and indigenous community knowledge.

## Acknowledgment

We thank the farming community of Balasore district, Odisha, for their participation and our field assistant for the support. We thank the Department of Zoology, the University of Calcutta for providing all the necessary requirements for the work. This work has been a part of research funded by Rufford Small Grant (Application Id. 18506-1)

## Authors’ contribution

Both the authors contributed to the study conception and design. Material preparation, data collection and analysis were performed by Deyatima Ghosh. The first draft of the manuscript was written by Deyatima Ghosh and both authors commented on previous versions of the manuscript. Both the authors read and approved the final manuscript. Concept-PB & DG, Data collection-DG, Analysis-DG & PB, Write-up-PB & DG

## Supporting Information

**S1 Figure** Schematic representation of applying the inter-reliability test between herpetofauna recognition checklist from each village and from the two validation groups

**S1 Table** Inter-reliability between data for herpetofauna recognition checklist from each village and the two validation groups

